# Dissecting antibiosis resistance to *Phthorimaea absoluta* in wild and cultivated tomato accessions

**DOI:** 10.64898/2026.07.11.737942

**Authors:** Komla Exonam Amegan, Florent Magot, Nicolas Desneux, Sylvia Del-Valle, Sylvia Salgon, Alan Kergunteuil, Bernard Caromel, Romain Larbat, Anne-Violette Lavoir

## Abstract

Tomato production faces a persistent challenge from the tomato leaf miner, *Phthorimaea absoluta*, a pest that severely limits yields while effective resistance in cultivated varieties remains scarce. To address this gap, wild tomato relatives represent a promising reservoir of resistance traits. In this study, 24 tomato accessions, including both cultivated types and wild species, were evaluated under greenhouse (no-choice) and tunnel (choice) conditions. Resistance mechanisms were characterized through measures of antibiosis such as leaflet lesion type, proportion of attacked leaflets, and mine density. The results revealed substantial variation between and within species, allowing classification of accessions into resistant, intermediate, and susceptible groups through multivariate analysis. Notably, the wild accession *Solanum habrochaites* ‘PI248707’ exhibited strong resistance, in contrast to susceptible cultivated varieties such as Rose de Berne. Under choice conditions, ‘PI248707’ sustained limited damage and disrupted larval development, with early instar larvae present but few reaching advanced stages, indicating an inhibitory defense response. Untargeted metabolomic profiling further highlighted pronounced constitutive differences between wild and cultivated accessions, with *S. pennellii* and *S. habrochaites* displaying higher metabolic diversity. By integrating phenotypic and metabolic data, specific metabolite classes associated with resistance were identified. These findings underscore the potential of wild tomato germplasm in breeding programs, with ‘PI248707’ standing out as a strong candidate for resistance introgression.

## 1 Introduction

The global spread of the tomato leaf miner, *Phthorimaea absoluta*, has emerged as a major constraint on tomato production, threatening yields in nearly all regions where the crop is cultivated (Biondi et al., 2018; Desneux et al., 2010). Native to the Peruvian highlands, this invasive lepidopteran has rapidly colonized multiple continents due to its high dispersal capacity, broad thermal tolerance, and rapid reproductive rate (Campos et al., 2017; Desneux et al., 2010, 2011; Machekano et al., 2018; Marchioro et al., 2017). Its larvae feeds internally on leaves, stems, buds, and fruits, causing extensive damage that can result in yield losses of up to 100% under severe infestations (Desneux et al., 2010; Gebremariam, 2015). This threat is exacerbated by the absence of effective resistance in the cultivated tomato.

Despite this vulnerability, tomato (*Solanum lycopersicum*) remains one of the most important vegetable crops worldwide, both nutritionally and economically (Gatahi, 2020; Qi et al., 2021). In 2022, global production reached 186 million tons across 5.17 million hectares (FAOSTAT, 2024), highlighting the scale at which this pest jeopardizes food production systems.

Management of *P. absoluta* has long relied on chemical insecticides. However, their intensive use has raised environmental and health concerns and has led to the rapid emergence of resistant pest populations (Roditakis et al., 2018). As a result, more sustainable strategies are being developed within Integrated Pest Management (IPM) frameworks (Desneux et al., 2022). Biological control approaches, including the use of egg parasitoids such as *Trichogramma spp*. and predators like *Nesidiocoris tenuis* and *Macrolophus pygmaeus*, have shown promising results, particularly under greenhouse conditions (Cherif et al., 2023; Cocco et al., 2013; Ferracini et al., 2019; Konan et al., 2021). Other alternatives, such as microbial agents and botanical insecticides, including essential oils, have also been investigated, although their effectiveness remains limited (Desneux et al., 2022; El-Aassar et al., 2015).

Within this context, the development of genetically resistant tomato varieties represents a critical and still unmet objective within sustainable management strategies for *P. absoluta* (Krokida et al., 2026). To date, no commercial cultivars exhibit effective resistance to *P. absoluta*. This limitation likely results from domestication bottlenecks that reduced the diversity of genes involved in plant defense, particularly those associated with the production of protective secondary metabolites (Oliveira et al., 2009; Salazar-Mendoza et al., 2023). In contrast, wild tomato relatives offer a valuable reservoir of genetic diversity that can be exploited to improve pest resistance. Species such as *Solanum habrochaites and Solanum pennellii* have been reported to exhibit strong resistance to *P. absoluta* (Bitew, 2018; de Azevedo et al., 2003; Maluf et al., 2010; Sridhar et al., 2019). Other closely related species, including *Solanum galapagense*, *Solanum cheesmaniae*, and *Solanum pimpinellifolium,* have demonstrated resistance to additional pests such as *Bemisia tabaci* and *Tetranychus urticae* (Rakha et al., 2017). Importantly, these species remain sufficiently compatible with cultivated tomato to be used in breeding programs aimed at introgressing resistance traits while minimizing undesirable agronomic effects (Mahmoud et al., 2022).

Resistance in wild tomato species is often associated with antibiosis, a mechanism by which plant traits negatively affect pest survival, development, or reproduction after host acceptance (Dias et al., 2019; Maluf et al., 2010). Antixenosis may also contribute by reducing host attractiveness or acceptance by herbivores (see (Amegan et al., 2024). Antibiosis is typically mediated by plant allelochemicals, such as phenolics and alkaloids, which interfere with insect physiology and feeding behavior (Ali et al., 2019; Painter, 1951; War et al., 2012). These biochemical defenses represent valuable targets for breeding programs seeking to enhance pest resistance through the selection and introgression of key metabolic traits.

In this study, we assessed antibiosis-based resistance to *P. absoluta* across 24 tomato accessions, including cultivated *S. lycopersicum* (large-fruited and cherry types) and three wild relatives (*S. habrochaites, S. pennellii*, and S*. cheesmaniae*). Cultivated accessions are typically classified as red-fruited, whereas *S. habrochaites* and *S. pennellii* belong to the green-fruited clade. We first screened for resistance using a no-choice assay, monitoring leaf damage over a 15-day period following infestation and quantifying multiple indicators to establish robust phenotyping criteria. We then validated these results under more realistic conditions by conducting a choice assay in tunnel conditions, comparing the most resistant and most susceptible accessions. Finally, we characterized the constitutive metabolic diversity of these accessions using untargeted LC-MS metabolomics. Comparative analyses between contrasted accessions were performed to identify metabolic signatures potentially associated with resistance mechanisms.

## 2 Material and methods

### 2.1. Plant materials and growth conditions

The accessions used in this study were taken from the diversity panel described in (Amegan et al., 2024, 2026), which represents a broad spectrum of genetic diversity within the tomato clade. These accessions were selected to include both cultivated and wild tomato lineages, spanning taxa with contrasting ecological adaptations (Moyle, 2008).

Seeds were pre-germinated in a controlled environment chamber at 24 ± 1°C, with 65 ± 5% humidity and a 16-hour light cycle. For *S. cheesmaniae*, seeds were previously soaked in bleach (2°4) for 30 min, to accelerate the degradation of the outer seed coat and, thereby, to promote germination. The germinated seedlings were initially grown in small cubic pots (7 × 7 × 6.5 cm) filled with a mixture of potting soil and perlite (1:1). After 10 days, the plants were transplanted into larger pots (12.8 × 12.8 × 20 cm) and moved to a greenhouse compartment regulated with a targeted temperature of 28°C during the day and 18°C at night. The greenhouse conditions allowed natural light variations according to the season, while humidity was maintained at approximately 70% during the experiment. The watering regime was adjusted to 20 L per greenhouse once daily (277 mL per plant) for both wild and cultivated tomato accessions.

### 2.2. Insect rearing and eggs production

A colony of *P. absoluta* was established from larvae collected from infested tomato plants at the INRAE Sophia Antipolis experimental station. They were reared in cages containing fresh tomato plants and adult moths were also fed with cotton wads soaked in a 50% honey-water solution to ensure proper nutrition. To collect eggs for the infestation experiments, newly emerged adult moths (100 males and 100 females) were placed in other rearing cages containing fresh tomato leaves supported by Oasis® floral foam. After six days, the eggs were collected using brushes and transferred onto leaflets of experimental plants in the early morning to ensure minimal handling damage.

### 2.3. Experimental design

#### 2.3.1. No-choice assays - Resistance phenotyping

The experiment was conducted in greenhouse conditions at the Institut Sophia Agrobiotech, INRAE, during the 2022 fall season. In the no-choice assay, each of the 24 accessions was replicated three times in two randomized blocks, resulting in a total of 144 plants (Fig. 1A). Each plant was inoculated with two eggs on the third leaf from the top, with one egg placed on the terminal leaflet and the other on an adjacent leaflet. The infested leaves were then covered with fine mesh nets to prevent the larvae from escaping (Fig. 1B, C). After egg hatching, the plants were monitored for 15 days, a period determined from previous experiments and the knowledge on *P. absoluta* life cycle to avoid adult flying escape risk.

**Figure 1:**
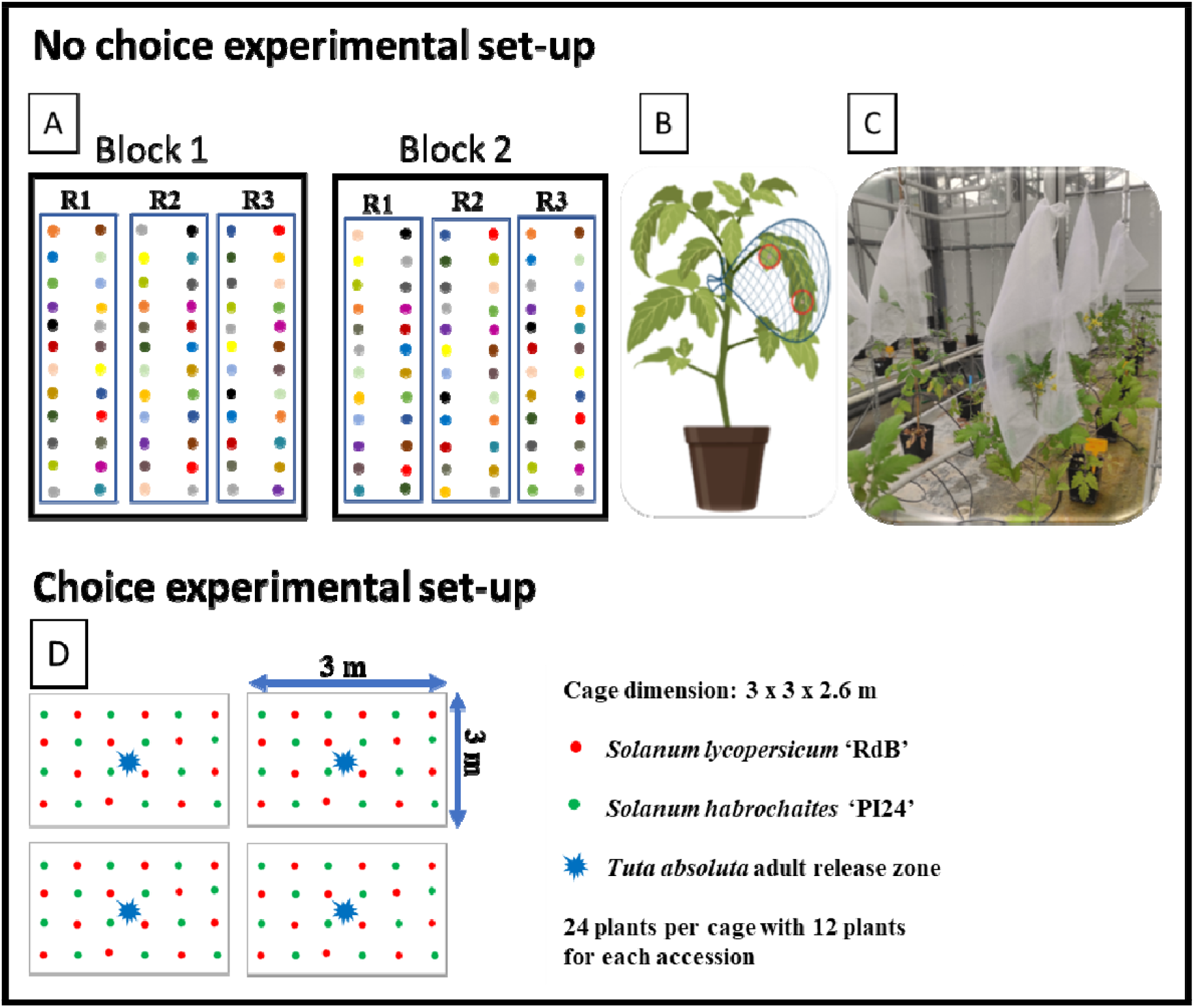
Experimental setups for choice and no-choice assays evaluating Tuta absoluta preferences on different Solanum accessions. A) No-choice experimental setup with two experimental blocks (Block 1 and Block 2), where colored dots represent 24 different Solanum accessions, each replicated three times per block (R1, R2, R3 = intra-block replicates); B) illustration of a single potted plant with the infested leaf covered using nylon mesh; C) photograph of actual greenhouse plants with mesh covers. D) Choice assay experimental setup diagram under tunnel conditions: red dot represents the cultivated variety ’RdB’ and green dot indicates the wild accession S. habrochaites ’PI24’. Cage dimensions: 3 × 3 × 2.6 m. Tuta absoluta adult release zone (blue star). 24 plants per cage with 12 plants for each accession.

To assess plant resistant phenotypes, three key phenotypic criteria based on plant damage were measured: (1) Leaflet Lesion Type (LLT), a visual score assessing the severity of leaf damage from 0 (no damage) to 5 (high damage); (2) the Percentage of Leaflets Attacked (PLA), assessed on a 0–5 scale, where 0 indicates no attacked leaflets and 5 corresponds to more than 80% of leaflets attacked (Dias et al., 2019; Maluf et al., 1997, 2010; Resende et al., 2006) and (3) the number of mines on attacked leaves, categorized as small (<0.5 cm) or large (>0.5 cm). Monitoring larval development proved to be impossible because the inoculated leaves were under the mesh, making observation of growth stages difficult

#### 2.3.2. Phenotypic validation in a choice assay

During the 2023 fall season, a choice assay was conducted in the tunnel at the Breeding Centre of Takii Seed in Eyragues, France (43.8508°N, 4.8297°E). This experiment aimed to compare the most resistant accession, *Solanum habrochaites* ‘PI248707’, identified from the no-choice assay, with the highly susceptible *S. lycopersicum* variety “Rose de Berne” (RdB). A total of 200 adult *P. absoluta* moths, with an equal male-to-female ratio, were released into large cages (3 × 3 × 2.6 m), spaced 1.5 meters apart and covered with insect-proof mesh (Fig. 1D). Each cage housed a mixed crop of 12 ‘PI248707’ plants and 12 ‘RdB’ plants arranged in a staggered pattern. The assay was replicated across four cages (n = 4 cages) (Fig. 1D). Three weeks after the moths were released, plant damages and pest development were recorded by measuring the total number of mines per plant, the total number of attacked leaves (TNLA), and the leaflet lesion type (LLT) score across the entire plant. We also recorded the number of *P. absoluta* larvae per plant, and the associated stages of larval development for each individual plant.

### 2.4. Tomato leaf metabolomic analysis

#### 2.4.1. Plant growth and extraction procedure

The constitutive metabolic content of leaves from the 24 accessions was assessed from non-infested plants grown in a climatic chamber under conditions described in Amegan et al. (2024). Two fresh leaflets from 4- to 6-week-old plants of each accession were harvested, immediately frozen in liquid nitrogen, and stored at −80 °C until freeze-drying. Metabolite extraction was performed following the procedure developed by (Royer et al., 2013) and reproduced in (Amegan et al., 2026).

#### 2.4.2. UHPLC-ESI-HRMS Analysis

UHPLC-ESI-HRMS data acquisition was performed as previously described in (Amegan et al., 2026). In brief, metabolite profiling was carried out using a Vanquish UHPLC system equipped with a binary pump, autosampler and temperature-controlled column, coupled to an Orbitrap ID-X mass spectrometer operating in both positive and negative electrospray ionization modes. Detailed chromatographic conditions, instrument parameters and acquisition settings are provided in the Data in Brief article (Amegan et al., 2026).

In the present study, the published raw dataset was reprocessed using an untargeted metabolomics workflow. Peak detection, chromatographic alignment and feature grouping were performed with Compound Discoverer 3.3 (Thermo Fisher Scientific). The full MS1–MS2 dataset was exported in Mascot Generic Format (.mgf) and analyzed with SIRIUS 5 for molecular formula annotation (SIRIUS and ZODIAC) and molecular structure prediction (CSI:FingerID), and compound classes were assigned using CANOPUS based on the NPClassifier ontology (Djoumbou Feunang et al., 2016; Dührkop et al., 2019; Hoffmann et al., 2021; Kim et al., 2021).

To construct the chemical class distribution presented in Figure 5, the NPClassifier ontology was adapted to improve readability. Carbohydrates were subdivided into “Acylsugars” and “Other carbohydrates”; alkaloids into “Polyamines” (ornithine alkaloids), “Glycoalkaloids” (pseudoalkaloids) and “Other alkaloids”; shikimate- and phenylpropanoid-derived compounds into “Flavonoids” and “Other phenylpropanoids”; and terpenoids into “Monoterpenoids”, “Sesquiterpenoids” and “Other terpenoids”. Final annotations were curated in Compound Discoverer 3.3 by combining elemental composition prediction with targeted searches in ChemSpider, HMDB and LipidMaps, and by inspecting spectral matches in mzCloud, MoNA and GNPS for the most relevant features.

### 2.5. Statistical analysis

To assess antibiosis resistance in the non-choice test, infestation levels were compared among the 24 tomato accessions belonging to 4 species. To reduce the impact of extreme values and ensure better statistical interpretation, a rank inverse normalization transformation (RINT) was applied, using the “orderNorm” function from the “bestNormalize” package (Peterson, 2021). This transformation allows homogenize the data while preserving their relative structure, thus ensuring better robustness of the analyses of variance. All statistical analyses were conducted on these transformed data. ANOVA was performed to assess the effects of accession and experimental block on all variables. Mean comparisons were then conducted using Fisher’s least significant difference (LSD) test, and homogeneous groups were identified with the LSD.test function from the *agricolae* package.

On the same set of data, Principal Component Analysis (PCA) was performed using the cluster, factoextra, and ade4 packages to reveal potential resistance profiles (Chessel et al., 2004; Kassambara & Mundt, 2016); Subsequently, Ward’s hierarchical method (Ward Jr, 1963) was applied for cluster analysis to classify the 24 accessions into distinct resistance groups. For the phenotypic data collected during the choice assays, all statistical analyses were also based on transformed data. ANOVA was performed, followed by pairwise mean comparisons using t-tests to identify significant differences between the resistant ‘PI248707’ accession and the susceptible ‘RdB’ accession. Additionally, Pearson correlation analysis was conducted to evaluate the relationship between larval development stages and infestation levels.

To investigate potential relationships between resistance and genetically driven phytochemical diversity, we used the phylogenetic tree previously established in (Amegan et al., 2024) and overlaid metabolomic information corresponding to 13 chemical classes involved in plant immunity. Multivariate analyses of metabolic composition were performed using permutational multivariate analysis of variance (PERMANOVA) based on Bray–Curtis dissimilarity matrices. Homogeneity of multivariate dispersion among species was assessed using the betadisper procedure followed by a permutation test. To identify the metabolic classes contributing most to the observed interspecific dissimilarity, a similarity percentage (SIMPER) analysis was performed on the relative abundance matrix using Bray–Curtis dissimilarities. All multivariate analyses were conducted with 9,999 permutations, and statistical significance was assessed at α = 0.05. More specifically, after identifying the most resistant accessions within *S. habrochaites* and *S. cheesmaniae*, enrichment analyses of metabolic classes were performed to identify metabolite categories that were significantly enriched or depleted relative to closely related highly susceptible accessions. All statistical tests were done using R software V4.3.1 (https://www.r-project.org/).

## 3. Results

### 3.1. No-choice experiments highlighted inter- and intraspecific diversity in resistance to *Phthorimaea absoluta*

The evaluated tomato accessions displayed a significant variation in their resistance to *P. absoluta* infestation according to estimates of LLT, PLA, number of small mines and large mines (p < 0.05) (Fig. 2, supplementary file S1, S2, S3). Globally LLT values were higher (meaning more damage) in *S. lycopersicum* and *S. pennellii* than in *S. cheesmaniae* and *S. habrochaites*. All *S. lycopersicum* were susceptible (LLT 3,5-5) when other families showed variability *i.e.* some accessions were susceptible when others were resistant (*S. pennellii* LLT 2,5-4; *S. cheesmaniae* LLT 1,5-4; *S. habrochaites* LLT 1-3,5). The accession ‘PI248707’ (*S. habrochaites*) was the least affected, whereas the *S. lycopersicum* ‘RdB’, ‘LA2078’ and ‘PI438’ together with the *S. cheesmaniae* ‘LA1450’ and the *S. pennellii ‘*LA1376’ exhibited the highest mean values (LLT >4) (Fig. 2). The PLA values followed the same general pattern as LLT to a lesser extent: Some accessions of *S. habrochaites* (PI248707, G1.1560, LA1033) and *S. cheesmaniae* (LA0746, LA0429) exhibited lower PLA values than *S. pennellii and S. lycopersicum* (supplementary file S1). Regarding the mines criteria, the *S. lycopersicum* accessions (LA2078, Maroc), and *S. cheesmaniae* (LA1450) showed a significantly lower mean value for small mines compared to the *S. cheesmaniae* accession LA0746 (supplementary file S2). In contrast, several accessions of *S. cheesmaniae, and S. habrochaites* exhibited fewer large mines, with no significant difference among many of them (supplementary file S3).

**Figure 2:**
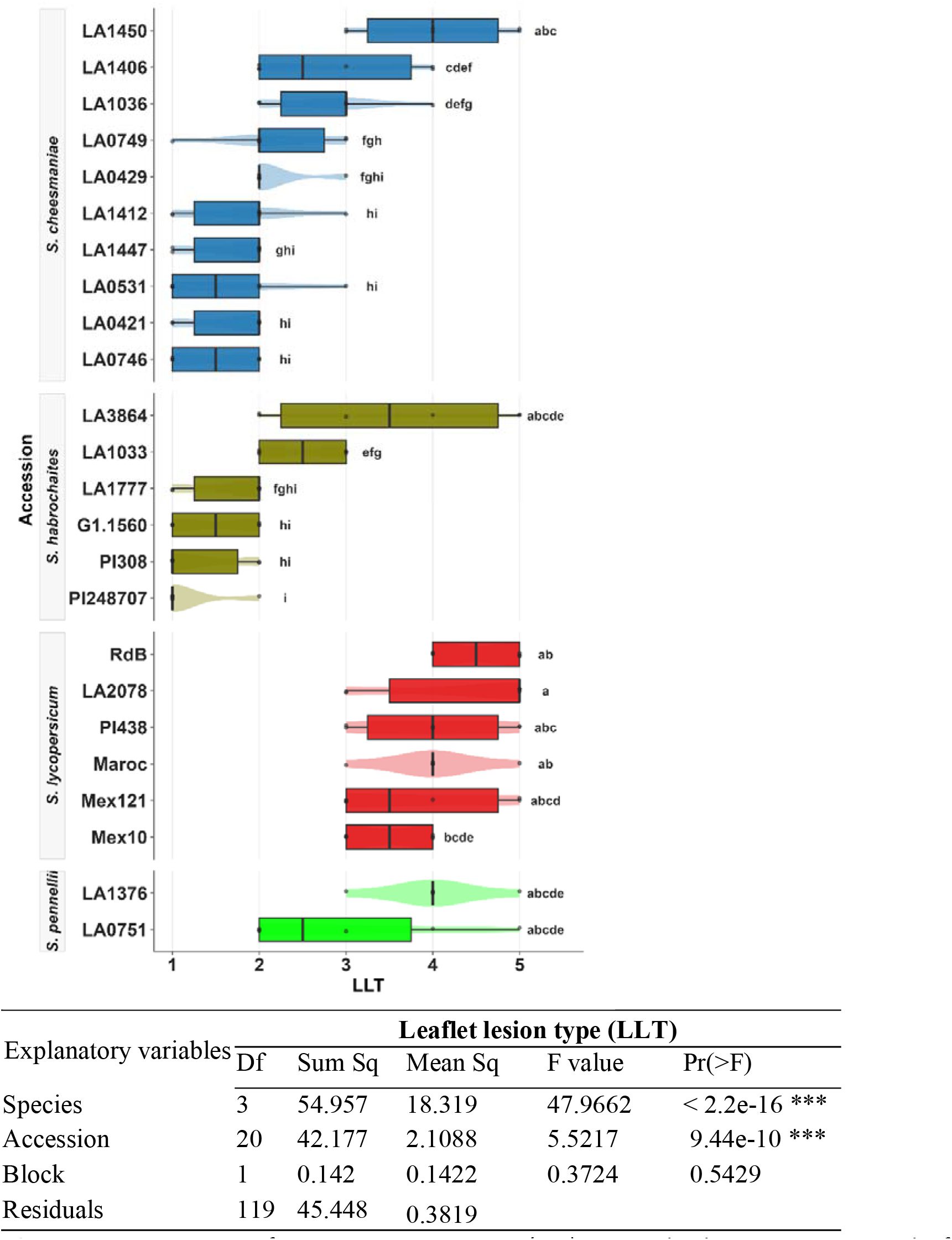
Comparison of resistance parameter (LLT) to T. absoluta across a panel of 24 Solanum accessions grown under greenhouse conditions in Sophia-Antipolis during the 2022 fall season. Different letters indicate significant differences using the LSD test at 5%. Mean value ± standard error (n=6). Mean values followed by a letter in common were not significantly different according to tukey test (p<0.05). LLT score: 0 = no lesion; 1 = small, rare lesions; 2 = small to medium-sized lesions, infrequent and often near the leaflet borders; 3 = medium to large lesions, coalescent with deformed leaflet borders; 4 = large, coalescent lesions with deformed leaflets; 5 = entire leaflet surface damaged. The results of the statistical test are given in the table below the figure.

Principal component analysis (PCA) on resistance parameters to *P. absoluta* revealed three distinct resistance/susceptibility groups, with the first two components explaining 87.5% of the total variance (Fig. 3). Axis 1 (60.5% of the variance) was strongly correlated with damage parameters (LLT, PLA, large mines), opposing highly resistant accessions (on the left, mainly *S. habrochaites, and S. cheesmaniae*) to susceptible accessions (on the right, *S. lycopersicum*). Cluster analysis supported this classification as the most resistant accessions in group 1 (Fig. 3, pink color) formed a distinct cluster from the most susceptible ones (LA1450, Maroc, RdB, and LA2078, (Fig. 3, yellow color)). Axis 2 (27%) highlighted differences among susceptible accessions based on the number of small mines, suggesting that some accessions allowed initial larval establishment but limited further development (more small mines but less large mines).

**Figure 3:**
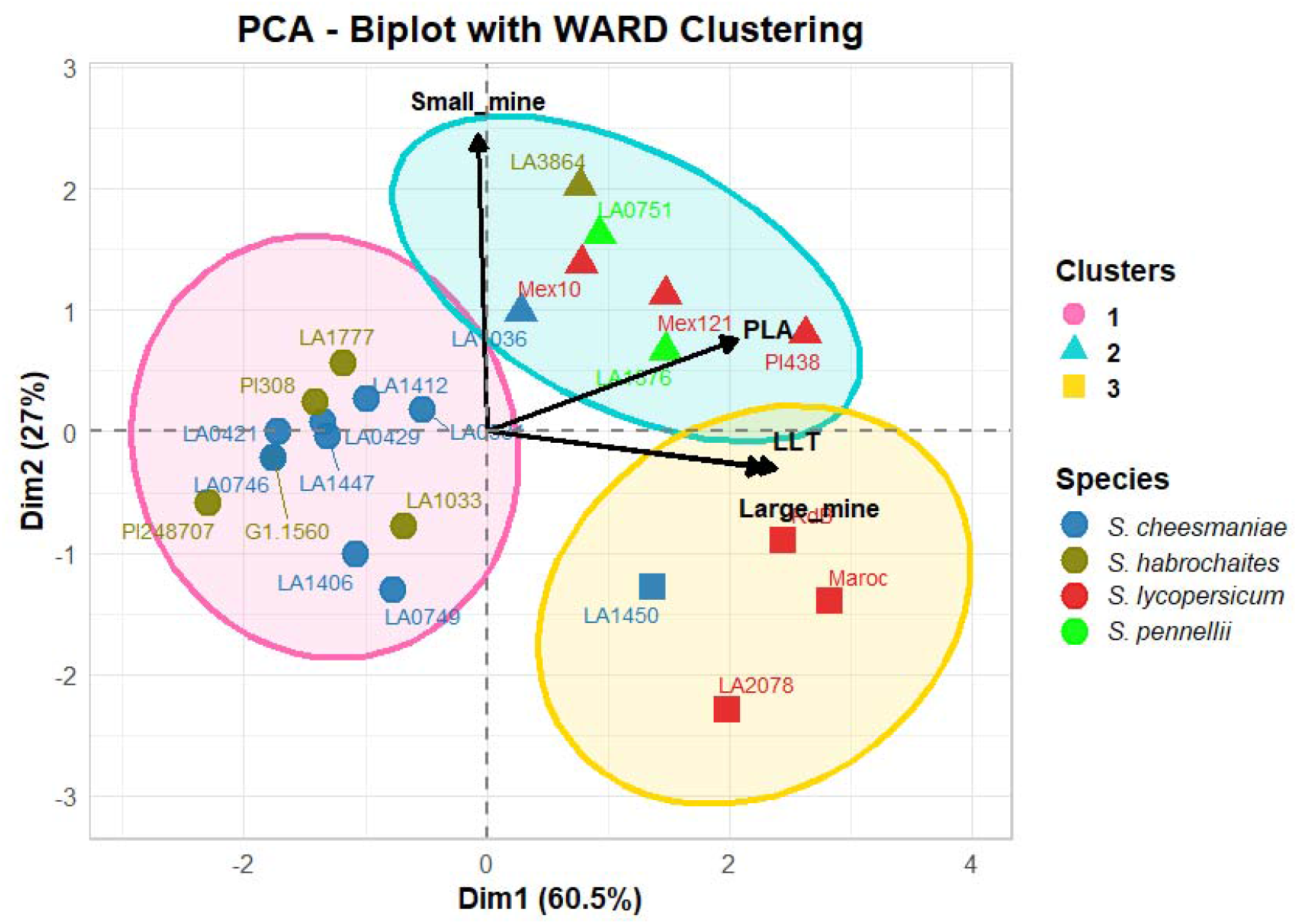
Principal component analysis (PCA) biplot with WARD clustering of tomato accessions based on Tuta absoluta resistance traits. The first two components explain 87.5% of the total variance (Dim1: 60.5%, Dim2: 27%). Arrows represent resistance variables: LLT (leaflet lesion type), PLA (percentage Leaflet Attacked), Small_mine and Large_mine (mining damage types). Cluster 1 = resistant accessions; Cluster 2 = susceptible with more small mines; Cluster 3 = susceptible with less small mines.

### 3.2. Choice assay results confirm the resistance of PI248707 to *Phthorimaea absoluta*

Based on the results of the no-choice experiment, the two accessions, PI248707’ and ‘RdB’, were selected for further evaluation in a choice assay to confirm the resistance of ‘PI248707’ to *P. absoluta*, compared to the susceptible ‘RdB’. The number of larvae at different developmental stages (L2, L3, L4) was significantly affected by the tomato accession (p < 0.05; Fig. 4A, B, C). Indeed, PI248707’ had a relatively higher number of larvae in the early L2 stage compared to ‘RdB’. However, the opposite trend was observed for later stage larvae (L3 and L4). As observed for the no choice assay, the mean value of LLT was significantly lower for ‘PI248707’ than for ‘RdB’, indicating that ‘PI248707’ sustained substantially less damage from *P. absoluta* (Fig. 4D). No statistically significant differences were observed between the two accessions for the total number of mines (No. mine) and the total number of leaves attacked (TNLA) (Fig. 4E-F).

**Figure 4:**
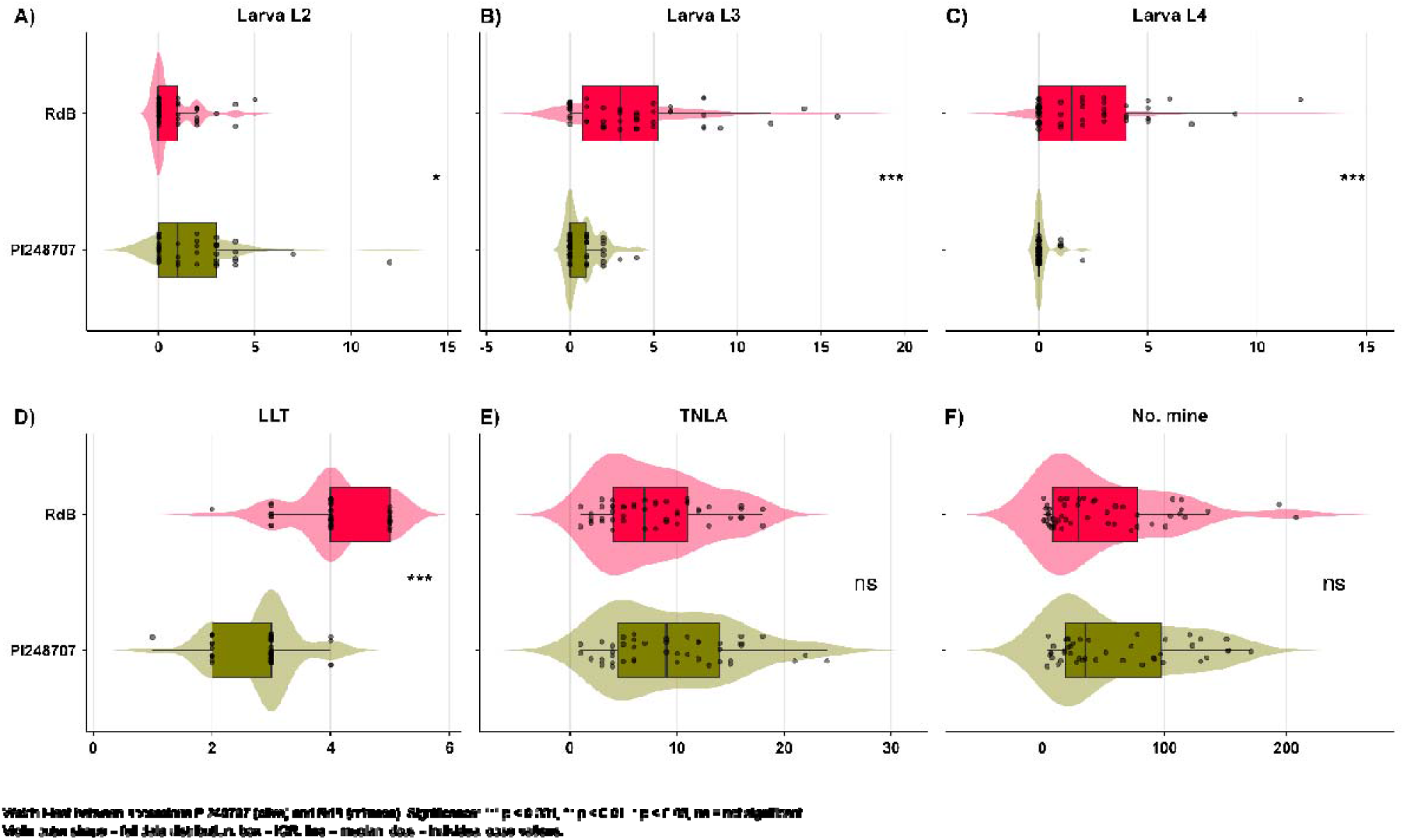
Tuta absoluta larva number at different developmental stages and Mean comparison of damage parameters. A) L2 instar larva (Larvae L2), B) L3 larva, C) L4 larvae, D) mean of Leaflet lesion type (LLT) on tomato plant lesion, E) mean of total number of leaves attacked (TNLA), F) total number of mines in the choice experiment comparing the wild accessions ‘PI248707’ with cultivated accession RdB using t.test at 5% for mean comparison.

**Figure 5:**
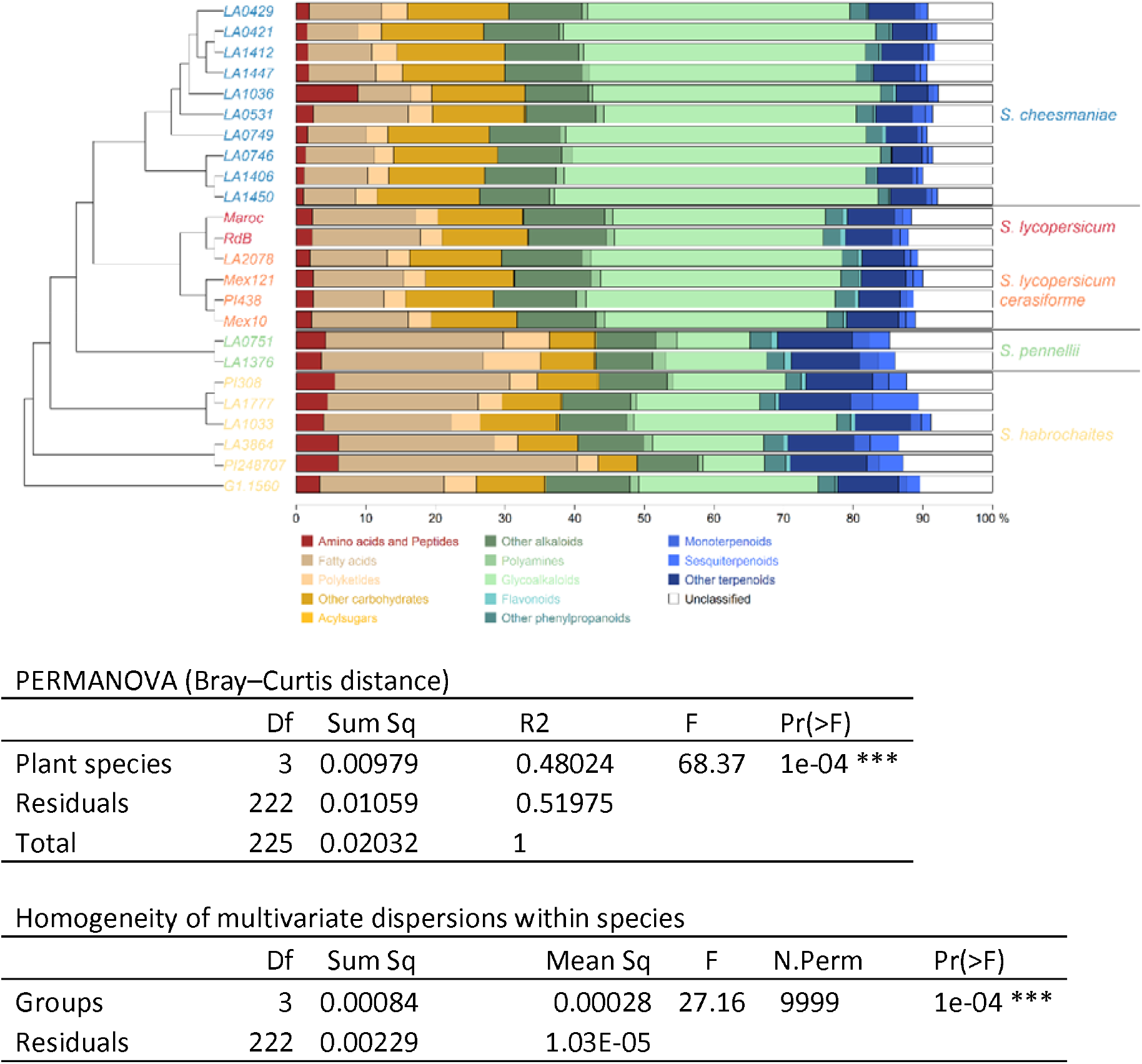
Tree-map representation of the relative abundance of metabolic classes in four Solanum species including S. lycopersicum (red-fruited), S. habrochaites and S. pennellii (green-fruited), and S. cheesmaniae. Metabolic features were assigned to metabolic classes using the CANOPUS software and the NP-Classifier pathway and superclass ontologies (Kim et al., 2021). PERMANOVA and multivariate dispersion analyses revealed significant differences in metabolite profiles among species, along with significant intraspecific variation, as shown in the tables below the figure.

Beyond these specific differences between accessions, correlation analyses revealed important relationships between the measured parameters. A strong positive correlation was found between the number of mines and the total number of leaves attacked (r = 0.83) (Table 1), indicating that leaf damage increases proportionally with infestation intensity, until it asymptotically approaches a plateau. The number of mines was also moderately correlated with the number of larvae at the L2 and L3 stages (r = 0.48 and r = 0.47, respectively), suggesting that early to mid-stage larvae contribute most to leaf mining. In contrast, the LLT score, representing leaf lesions, was moderately correlated with L3 and L4 larvae (r = 0.53 and r = 0.63), reflecting cumulative damage over time. Overall, these results highlight a stronger association between early larval stages and mining activity, while later stages may have the strongest impact in terms of largest mines and damage.

**Table 1:**
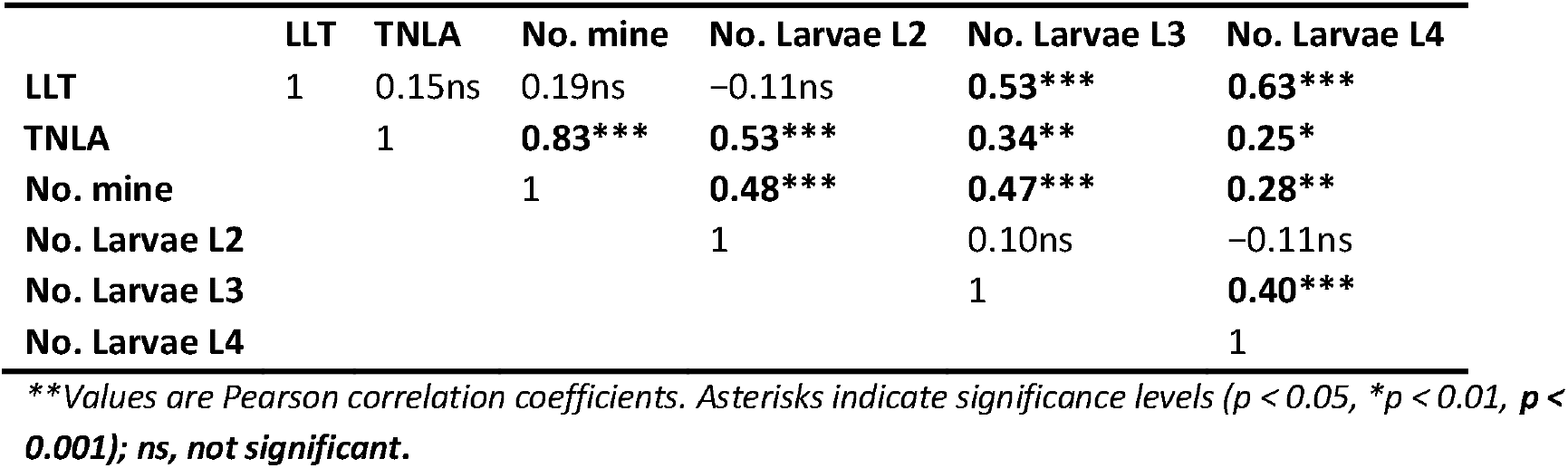
Pearson correlation matrix among leaf lesion type (LLT), total number of leaflets attacked (TNLA), number of mines, and larval stages (L2–L4) recorded in the choice assay.

### 3.3. Clade-wide chemodiversity of constitutive foliar metabolic composition

Untargeted LC-MS analysis of hydro-methanolic leaf extracts from 24 tomato accessions generated 3,764 metabolic features (Amegan et al., 2026). This dataset revealed a strong phylogenetic structuring of constitutive foliar metabolic composition, with distinct biochemical profiles among cultivated tomato and its wild relatives (Fig. 5).

Metabolic profiles differed significantly among species (PERMANOVA: F₃,₂₂₂ = 68.37, R² = 0.48, *p < 0.001*), indicating that species identity explained 48% of the variation in constitutive chemical composition. Wild species, namely *S. pennellii* and *S. habrochaites*, were characterized by higher relative proportions of amino acids and peptides, fatty acids, and terpenoids (including mono- and sesquiterpenoids). They also contained a greater proportion of acyl sugars than *S. lycopersicum*, although this class represented only a small fraction of the total metabolome. Conversely, these two wild species exhibited lower proportions of carbohydrates and glycoalkaloids than *S. lycopersicum*. Overall, these compositional differences clearly reflect the well-established metabolic divergence between green-fruited and red-fruited tomato lineages.

Within-species metabolic variability also differed significantly among species (betadisper: **F**₃,₂₂₂ = 27.17, *p* < 0.001; Supplementary Table S5A-B). The two green-fruited species exhibited greater within-species metabolic heterogeneity than the red-fruited groups, with *S. habrochaites* showing pronounced variation in glycoalkaloid abundance, ranging from 8.9% in PI248707 to 29.1% in LA1033. In contrast, *S. lycopersicum* displayed comparatively homogeneous metabolic profiles consistently dominated by glycoalkaloids (32.5%), fatty acids (13.5%), and carbohydrates (12.4%). Consistent with this pattern, average metabolic profiles indicated a closer similarity between the two red-fruited species (*S. lycopersicum* and *S. cheesmaniae*) than with the green-fruited taxa (Supplementary Table S6).

SIMPER analysis further indicated that the metabolic differentiation among species was primarily explained by other alkaloids (32.9%) and fatty acids (22.8%), whereas glycoalkaloids contributed only marginally (4.0%) to the overall interspecific dissimilarity (Supplementary Table S7).

### 3.4. Pairwise differences of metabolites in contrast tomato accessions

Based on resistance data showing inter- and intra-specific variability within the accession panel, the constitutive metabolic composition of leaves from the most resistant (*R*) accessions was compared to that of the most susceptible (*S*). We first compared the metabolome of the most resistant accession ‘PI248707’ with the susceptible ‘RdB’ which does not belong to the same species. Then, inside *S. cheesmaniae* and *S. habrochaites,* we compared the most resistant and the most susceptible accessions.

At the inter-specific level, comparison of the leaf metabolome between PI248707 (*R*) and RdB (*S*) revealed a total of 2628 differential features, with 1560 over-accumulated in PI248707 and 1063 in RdB (supplementary file S4). Among them, enrichment analysis based on CANOPUS chemical class affiliation highlighted polyamines, as the most significantly enriched metabolic class in the resistant accession ‘PI248707’, followed by fatty acids, sesquiterpenoids, and other terpenoids. Conversely, the susceptible accession RdB showed significant enrichment in other carbohydrates and alkaloids (Fig. 6A).

**Figure 6.**
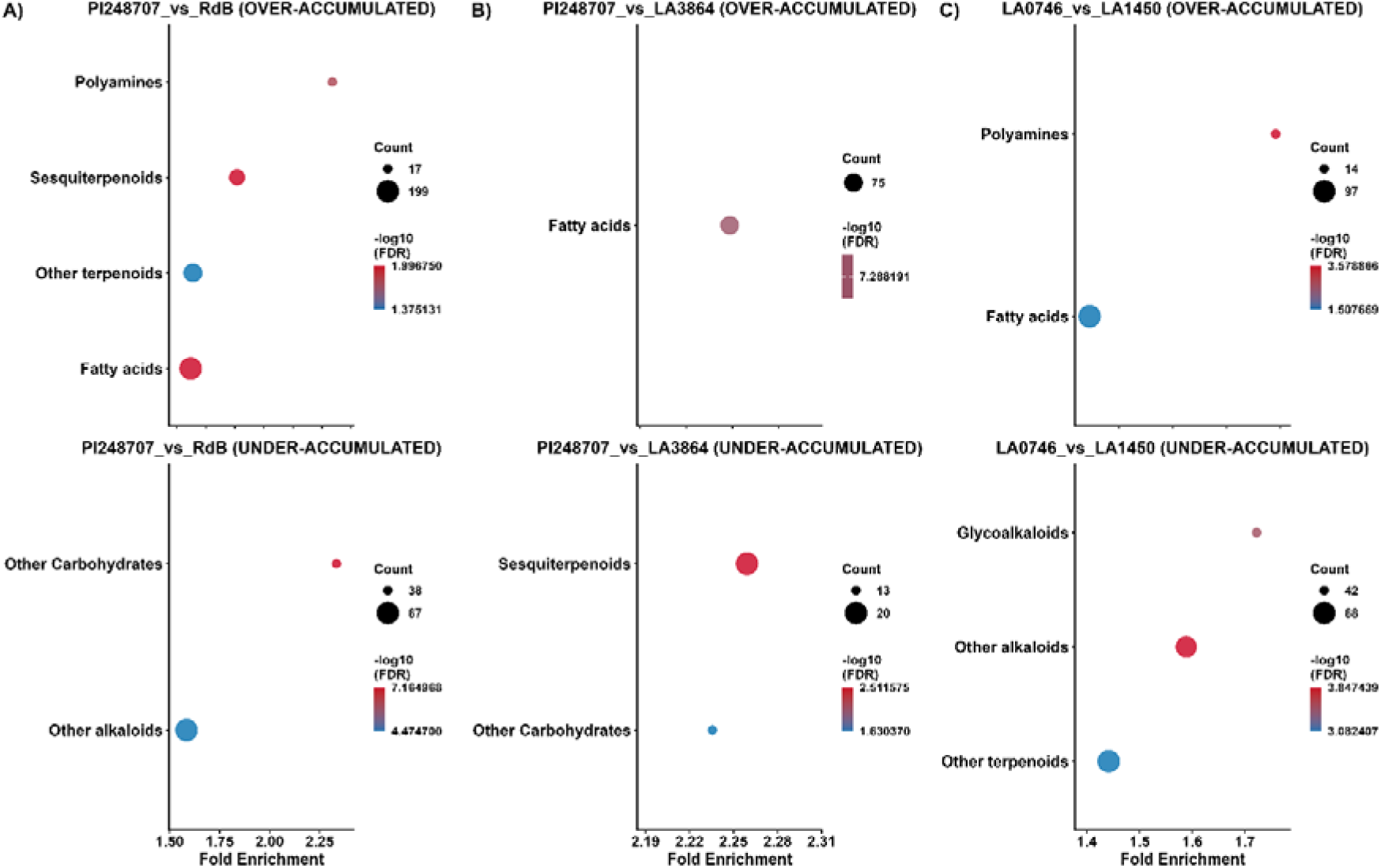
: Metabolic pathway enrichment in resistant vs. susceptible genotypes. (A) PI248707 vs. RdB, (B) PI248707 vs. LA3864, (C) LA0746 vs. LA1450. Bubble size indicates the count of significantly different metabolites, x-axis shows fold enrichment, and color intensity represents statistical significance (-log10(FDR) < 0.05). Top: upregulated pathways; Bottom: downregulated pathways in resistant compared to susceptible genotypes.

At the intra-specific level, distinct metabolic patterns were also detected between resistant and susceptible accessions. In *S. habrochaites*, comparison between PI248707 (R) and LA3864 (S) identified 751 significant features (363 up in PI248707 and 388 in LA3864), with enrichment analysis showing fatty acids as the most enriched class in the resistant accession, while the susceptible accession was enriched in sesquiterpenoids and other carbohydrates (Fig. 6B). In *S. cheesmaniae*, comparison between LA0746 (R) and LA1450 (S) revealed 1657 significant features (882 up in LA0746 and 775 in LA1450), with highly significant enrichment of polyamines in the resistant accession, contrasting with enrichment in alkaloids and glycoalkaloids in the susceptible accession (Fig. 6C). Overall, resistant accessions are characterized by enrichment in lipid-derived compounds (mainly fatty acids) and polyamines, whereas susceptible accessions preferentially accumulate alkaloids and carbohydrate-related metabolites.

## 4. Discussion

This study assessed resistance to *Phthorimaea absoluta* in a diverse set of wild and domesticated *Solanum* accessions under both no-choice and choice conditions. We identified robust damage-based indicators such as leaflet lesion type (LLT), percentage of leaflet attacked (PLA), and mine size and number that reliably distinguished resistant from susceptible accessions. Resistance varied markedly among and within species, with *S. habrochaites* and *S. cheesmaniae* showing the highest average resistance, although each species also included accessions with susceptibility levels comparable to *S. lycopersicum*. By integrating metabolomic and resistance data, we further revealed constitutive biochemical differences among species, with resistant genotypes being associated with defensive pathways involving fatty acids, polyamines, and terpenoids, whereas susceptible accessions showed a higher relative abundance of glycoalkaloids and carbohydrates

### 4.1. Assessing resistant accessions based on relevant resistance indicators

#### 4.1.1 Relevant indicators to assess tomato plant resistance against P. absoluta

Still, our results demonstrate that resistance to *P. absoluta* can be effectively characterized through a targeted set of phenotypic parameters. Resistance indicators, only based on plant damage (including leaf lesion type (LLT), percentage leaflet attacked (PLA) and the number of both small and large mines), successfully captured resistance diversity among accessions. PCA representation confirmed the reliability of these indicators, as resistant and susceptible accessions clustered distinctly (Fig. 4). LLT, which likely reflects feeding deterrence, was particularly informative, as a lower LLT score suggests higher antibiosis resistance, making it a valuable indicator (Dias et al., 2019; Mahmoud et al., 2022; Resende et al., 2006). The high informativeness of LLT is probably related to the biology of *P. absoluta*, a leafminer whose feeding activity directly impacts leaf tissue thickness and integrity. Unlike other pests that cause superficial or temporary damage, *P. absoluta* larvae create permanent structural changes in leaf architecture, making LLT a reliable indicator of cumulative damage severity. The applicability of this situation diminishes for pests whose damage is less visible or more ephemeral.

Also, the high correlations between PLA, number of large mines, and LLT in the no-choice test, alongside strong correlations between mine number and leaves attacked in the choice test, indicate measurement redundancy in our current protocol. However, the number of small and large mines alone does not fully capture resistance mechanisms. Our correlation analyses revealed significant relationships between LLT scores and mining patterns, suggesting that leaf structural characteristics may influence larval behavior. Specifically, higher LLT values (indicating more severe lesions) were associated with different mining patterns, as thicker or more damaged leaves could potentially impede larval development and mines establishment. A high number of small mines may indicate larval difficulty in finding a suitable feeding site rather than increased plant susceptibility (Leite et al., 2001). The negative correlation between small and large mines suggests that larvae may be abandoning feeding sites or experiencing developmental impairments that prevent them from reaching later instars.

For efficient resistance screening, we recommend focusing primarily on accurate LLT assessment, which most effectively captures damage severity across both assay types. When using this visual scoring method, standardized training and multiple independent evaluations are essential to minimize observer bias and prevent over an underestimation of resistance levels. This approach should be complemented by larval stage evaluation, which reveals early resistance mechanisms that impair larval development progression. It is also relevant to note that the variability in egg hatching rates across accessions could have influenced resistance assessments. Differences in leaf surface properties, micro-climate conditions, or chemical compositions might affect egg viability independently of actual plant resistance to larval feeding (Blackmer et al., 2002; Fatouros et al., 2016; Rojas et al., 2017). This represents an important confounding factor in our resistance evaluations. This streamlined yet precise approach maintains phenotyping accuracy while significantly reducing evaluation time and resources, enabling higher throughput screening of germplasm collections for resistance breeding programs.

#### 4.1.2 Assessing resistant versus susceptible accessions

Differences between wild and cultivated accessions were evident in the number of large mines, with lower values observed in wild species (Ecole et al., 2001, 2000; Suinaga et al., 2004). This suggests breeding potential from wild accessions, as fewer large mines indicate reduced pest burden.

For the choice experiment, we used the same plant resistance indicators but added data on pests and we confirmed the resistance of *S. habrochaites* ‘PI248707’ compared to the susceptible *S. lycopersicum* ‘RdB’ under natural host-selection conditions. Although the total number of mines and attacked leaves did not differ significantly between accessions, larval progression was markedly impaired on ‘PI248707’. This suggests an antibiosis mechanism where *P. absoluta* larvae face physical and/or chemical barriers preventing development beyond the L2 stage. Accession ‘PI248707’ defense restricted pest feeding and establishment, evidenced by reduced larval development at later stages (L3 and L4) (Fig. 4).

These findings support previous studies reporting that pest feeding and development is impaired on *S. habrochaites* accessions (Bitew, 2018; Mahmoud et al., 2022; Maluf et al., 2010). PI248707’s ability to hinder early larval development significantly reduces mine formation and foliar damage. The link between mine density and lesion severity emphasizes the importance of resistance mechanisms acting at the earliest larval stages. By preventing larvae from reaching advanced stages, ‘PI248707’ limits damage severity. In contrast, ‘RdB’ sustains greater foliar deterioration at later stages, confirming its susceptibility. These findings highlight the critical role of early-stage resistance in mitigating *P. absoluta* impact and offer valuable insights for breeding pest-resistant tomato cultivars.

While the ‘PI248707’ case study demonstrated clear resistance mechanisms, broader analysis across the entire accession panel revealed more complex patterns. Interestingly, antibiosis assays did not lead to such a clear pattern distinguishing the wild green fruited accessions as the more resistant and the domesticated one as susceptible. Indeed, although *S. habrochaites* accessions displayed more resistance, *S. pennellii* ones were as susceptible as the *S. lycopersicum* cultivars, and inversely *S. cheesmaniae* accessions which harbored a metabolic pattern close to *S. lycopersicum*, exhibited globally a resistance comparable to *S. habrochaites*. This result supports previous findings showing that *S. habrochaites* and *S. cheesmaniae* harbor resistance traits against *P. absoluta* (Bitew, 2018; Maluf et al., 2010; Rakha et al., 2017; Vitta et al., 2016). However, the susceptibility observed in *S. pennellii* accessions (LA0751, LA1376) contrasts with reports of strong resistance in this species (Bitew, 2018), suggesting intra-specific variability (Gharekhani & Salek-Ebrahimi, 2014). In the case of *S. pennellii,* only two accessions were evaluated in the present work, yet the observed contrast already points to the intra-specific diversity that becomes evident when a larger number of accessions are screened. Indeed, the significant intra-specific variation observed in *S. habrochaites* and *S. cheesmaniae* demonstrated that resistance to *P. absoluta* was not uniformly distributed within species. While most wild accessions showed strong resistance, others (e.g. LA3864 and LA1450), displayed susceptibility comparable to cultivated tomatoes, suggesting that defense metabolite accumulation varies considerably among accessions of the same species. While emblematic genotypes such as LA1777 in *S. habrochaites* and LA0716 in *S. pennellii* have been extensively studied as sources of resistance in plant insect interaction, this focus may have overshadowed the substantial intra-specific diversity that exists within wild species. Our findings revealed that resistance is not a fixed trait at the species level but rather varies significantly among accessions, emphasizing the need to broaden genetic exploration beyond a few well-characterized genotypes.

### 4.2. The constitutive metabolic diversity of *Solanum* species highlights several potential resistant markers

Linking leaf metabolic diversity to resistance traits through both inter- and intra-specific approaches has led to identifying candidate resistance biomarkers. This multi-level taxonomic framework allows for comprehensive assessment of chemical diversity patterns that may underline species-specific resistance mechanisms.

Untargeted LC-MS analysis of hydro-methanolic leaf extracts revealed 3,764 metabolic features across 24 tomato accessions, demonstrating distinct constitutive chemical profiles between wild green fruited species (*S. pennellii* and *S. habrochaites*) and cultivated species. Metabolite annotations further characterized these differences, confirming that wild tomato species display a higher proportion of compounds usually associated with plant defense, including amino acids/peptides, fatty acids, acyl sugars and terpenoids.

To go further, pathway enrichment analysis across the three resistance-contrasting comparisons consistently revealed the over-representation of well-established defensive metabolic pathways, particularly fatty acids, polyamines, and terpenoids, in resistant genotypes. These compounds are known to interfere with insect development through multiple mechanisms. For instance, polyamines exert cytotoxic effects on pathogen cells and induce oxidative stress damage in herbivores by disrupting oxidative stress signaling pathways (Kaur et al., 2010). Similarly, terpenoids compounds demonstrate repellent and antifeedant activities against *P. absoluta*, disrupting both feeding and oviposition behavior (Anastasaki et al., 2018; Kortbeek et al., 2023; Mahmoud et al., 2022).

Fatty acids likewise contribute significantly to plant defense through their dual role as signaling molecules and direct defensive compounds, notably known for their insecticidal properties (Boncan et al., 2020; Kumaraswamy et al., 2025; Lim et al., 2017; Plata-Rueda et al., 2018; Ramos-López et al., 2012; Walley et al., 2013). Unsaturated fatty acids like linolenic and linoleic acids serve as precursors to jasmonic acid and other oxylipins that orchestrate defense responses against herbivores (Browse, 2005; Farmer et al., 2003). Beyond signaling, fatty acids reinforce physical barriers through cuticular components while simultaneously exerting toxic effects on insect metabolism (Yeats & Rose, 2013). The elevated fatty acid levels observed in resistant *S. habrochaites* ‘PI248707’ might explain its enhanced protection against *P. absoluta*. These lipids not only strengthen structural defense but also activate systemic immune responses that prepare healthy tissues for potential threats (Yu et al., 2013).

In addition, the pair comparison revealed a contrasting pattern with the constitutive higher accumulation of many glycoalkaloid features in the leaf content of more susceptible accessions (Fig. 6). The high glycoalkaloid (GA) content in cultivated *S. lycopersicum* and its close relatives (*S. cheesmaniae*) compared to more distantly related wild species such as *S. habrochaites* and *S. pennellii* suggests that these compounds play a significant role in Solanaceae metabolism. However, despite their presumed toxicity, they do not seem to confer resistance against *P. absoluta*, as *S. lycopersicum* remains highly susceptible to this pest. This suggests that *P. absoluta*, as a Solanaceae specialist, may have evolved strategies to cope with glycoalkaloids, either by detoxification (e.g. cytochrome P450s, esterases, glutathione S-transferases, as widely documented in insecticide resistance) or by sequestration mechanisms that neutralize their toxicity (Amezian et al., 2021; Hilliou et al., 2021; Leite Dias & D’Auria, 2025; Lu et al., 2021). Such adaptations would parallel cases in other Solanaceae specialists, such as *Tequus sp*., a wild potato herbivore able to thrive on *S. commersonii*, which is rich in glycoalkaloids, while generalists avoid it (Altesor et al., 2014). The capacity of *P. absoluta* to metabolize or tolerate these compounds may thus represent a key adaptation to its host range. Additionally, glycoalkaloids are considered non-toxic storage forms of alkaloids, allowing plants to maintain defensive compounds in a “safe” state (glycosylated vs aglycone forms) until needed (McKey, 1974). However, this storage form might simultaneously facilitate their accumulation in tissues without impairing herbivores that have developed efficient detoxification systems.

These baseline metabolic differences between accessions demonstrate how species divergence shapes inherent defensive chemistry and provide crucial insights into the biochemical basis of *P. absoluta* resistance that could guide targeted breeding programs. Yet, constitutive chemical defenses only provide the primary plant defense barrier, while induced defenses are triggered upon pest recognition and constitute a secondary barrier. Validating the implication of these metabolites in the tomato resistance to *P. absoluta* would require additional experiments, such as temporal comparative analyses tracking metabolomic profile changes in resistant versus susceptible accessions before and after *P. absoluta* infestation. Combined transcripto-metabolomic studies are also required to establish direct correlations between defense gene expression and bioactive compound production.

## Conclusion

This study suggests the potentially important role of wild *Solanum* species, with *S. habrochaites* ‘PI248707’ emerging as a promising genetic resource for enhancing tomato resistance against *P. absoluta*. The genetic and biochemical diversity we identified opens promising avenues for developing naturally pest-resistant cultivars. While our findings highlight the potential of constitutive metabolites in conferring resistance, we acknowledge that our focus on these compounds does not capture inducible responses triggered upon herbivore attack. Some wild species may rely more heavily on induced rather than constitutive defense, representing an aspect not fully explored in our dataset. Future research directions should include validating candidate metabolic markers, identifying the genetic loci underlying this resistance, and optimizing their introgression into commercial tomato lines. If successfully integrated into plant breeding strategies, these discoveries could contribute to developing tomato cropping systems less reliant on insecticides against *P. absoluta*. The conservation of major metabolomic families across species, with primarily quantitative variations, raises an intriguing question: whether breeding programs might successfully enhance these constitutive defense mechanisms without compromising agronomic performance. Successfully addressing this intersection between natural defense and agricultural may constitute one of the most compelling challenges for the future of crop improvement. From a societal perspective, such research agenda about sustainable development of cultivars adapted to low-input systems may help to support agroecological transition and famer empowerment.

## Acknowledgements

We thank the Biological Resources Center of Avignon (CRB-Lég team of INRAE GAFL) for providing tomato seeds, the Plant Experimental Platform in Lorraine (PEPLor, University of Lorraine) for plant cultivation, the Metabolomic and Structural Analytic Platform (PASM, University of Lorraine) for metabolomic analyses, and Takii Seed for hosting the choice experiment as a breeding programme partner. We also thank the ISA technical service and greenhouse management teams, as well as Philippe Bearez and Lionel Salvy, for their technical support. We are grateful to Christophe Robin for his advice and for critically reading the manuscript, and to François Massol for his guidance on statistical analyses.

## Funding

This research was funded by the “CapZéroPhyto” project of the French Priority Research Programme “Cultiver et Protéger Autrement” (ANR grant number ANR-20-PCPA-0003).

## Conflict of interest disclosure

The authors declare no competing interests

## Data, scripts, code, and supplementary information availability

Data, R scripts, supplementary materials and Readme text are available online: Amegan, K., Magot, F., Desneux, N., Del-Valle, S., Salgon, Sylvia., Kergunteuil, A., Caromel, B., Larbat, R., & Lavoir, A.-V. (2026). Phenotypic, metabolomic and phylogenetic datasets with R workflows for the analysis of tomato resistance to Tuta absoluta [Data set]. In Peer Community in Plant Biology (Version v2). Zenodo. https://doi.org/10.5281/zenodo.21280659

## Notes

https://doi.org/10.5281/zenodo.21280659

### Competing Interest Statement

The authors have declared no competing interest.

